# Energy Deficit is a Key Driver of Sleep Homeostasis

**DOI:** 10.1101/2024.05.30.596666

**Authors:** Scarlet J. Park, Keith R. Murphy, William W. Ja

## Abstract

Sleep and feeding are vital homeostatic behaviors, and disruptions in either can result in substantial metabolic consequences. Distinct neuronal manipulations in *Drosophila* can dissociate sleep loss from subsequent homeostatic rebound, offering an optimal platform to examine the precise interplay between these fundamental behaviors. Here, we investigate concomitant changes in sleep and food intake in individual animals, as well as respiratory metabolic expenditure, that accompany behavioral and genetic manipulations that induce sleep loss in *Drosophila melanogaster*. We find that sleep disruptions resulting in energy deficit through increased metabolic expenditure and manifested as increased food intake were consistently followed by rebound sleep. In contrast, “soft” sleep loss, which does not induce rebound sleep, is not accompanied by increased metabolism and food intake. Our results demonstrate that homeostatic sleep rebound is linked to energy deficit accrued during sleep loss. Collectively, these findings support the notion that sleep functions to conserve energy and highlight the need to examine the effects of metabolic therapeutics on sleep.

## Introduction

Excessive eating following a sleepless night is a prevalent phenomenon in modern society^1,2^. Sleep and eating—typically mutually exclusive behaviors that are amongst those most vital for survival and prosperity—are intricately linked. Food deprivation increases activity levels in many species, including flies^3^, whereas food ingestion temporarily elevates immediate sleepiness—a phenomenon colloquially known as “food coma”—in flies^4^, laboratory rodents^5,6^, and humans^7,8^. Conversely, acute and chronic sleep deprivation is associated with altered taste perception, food cravings, increased appetite and food intake, rapid weight gain, and detrimental metabolic changes in laboratory rodents^9^ and humans^10^.

Sleep homeostasis has been hypothesized to serve as an adaptive response to energy expenditure, purposed towards reducing and recovering from energy use^11^. This is consistent with findings that the intensity of neuronal activity during wakefulness may contribute to sleep drive^12^. This may also explain the greater sleep need of nematodes during energetically expensive molting periods^13^. In at least some mammals, brown fat metabolism is necessary for sleep rebound following deprivation, further supporting the idea that energy expenditure generates sleep-promoting signals^14^. If this model is correct, then “hard” sleep loss—i.e. homeostatic sleep loss that accompanies a subsequent, compensatory increase in sleep (as opposed to “soft” sleep loss, which is not succeeded by sleep rebound)—should be associated with increased metabolic activity and/or hunger.

Sleep curtailment has been consistently linked to increased food intake^1,15,16^ and reduced energy expenditure^17^. However, understanding the precise mechanisms that drive changes in feeding and metabolism, and that regulate the sequence of events following sleep disruption, remains challenging due to the complex relationship between these factors. Experimental isolation of specific variables is particularly difficult in observational human sleep studies. Development of an effective model system for controlled manipulations and measurements of sleep and feeding would enable research on the regulatory pathways linking these processes.

Studies on the characteristics of sleep and its genetic and neuronal control have yielded diverse methods to manipulate sleep in *Drosophila melanogaster*^18,19^. These tools, combined with the scalability and genetic tractability of the model, provide unique opportunities for studying homeostatic sleep regulation and the underlying circuit logic^19,20^. Despite recent advances in understanding the interaction between sleep regulation and other homeostatic processes in flies, including metabolism and food intake^4,21^, studies on the relationship between sleep and feeding behavior have typically relied on measurements of each behavior in separate groups of animals, mainly due to the lack of techniques that enable paired measurements. We have previously shown that flies exhibit postprandial sleep in the Activity Recording CAFE (ARC), an assay that allows concurrent, longitudinal measurements of sleep and food intake in freely moving animals^4^. The ARC can thus account for individual variability, making correlational studies more powerful and offering more in-depth insights into the relationship between short-, intermediate-, and long-term food intake and sleep.

To investigate the link between sleep deprivation, energy balance, and sleep rebound, we measured changes in sleep, feeding, and metabolic expenditure with different methods of sleep restriction, including starvation-induced sleep suppression and thermogenetic stimulation of various wake-promoting neurons. Manipulations that elicit sleep rebound are accompanied by increased food intake during or following sleep deprivation, as well as increased respiratory metabolism—two dominant variables in the energy balance equation. These results suggest that homeostatic regulation of sleep is a result or a readout of energy balance.

## Results

We first tested the effect of mechanical sleep deprivation (SD) on food intake using a custom-made fly vial agitator (Supplemental Movie 1), which effectively prevented sleep as indicated by the subsequent sleep rebound observed in the Activity Recording CAFE (ARC), an apparatus that allows simultaneous measurements of movement and food intake of individual animals^22^ (Fig. 1a). Additionally, sleep-deprived flies consumed more food the following day despite the homeostatic increase in sleep, which correspondingly limits dining hours (Fig. 1a). Importantly, increased consumption was not indirectly due to compromised feeding during the deprivation treatment, as food intake was not significantly different during SD (Fig. 1b), confirmed by quantifying consumption of a radiolabeled medium during and after SD.

**Figure 1.**
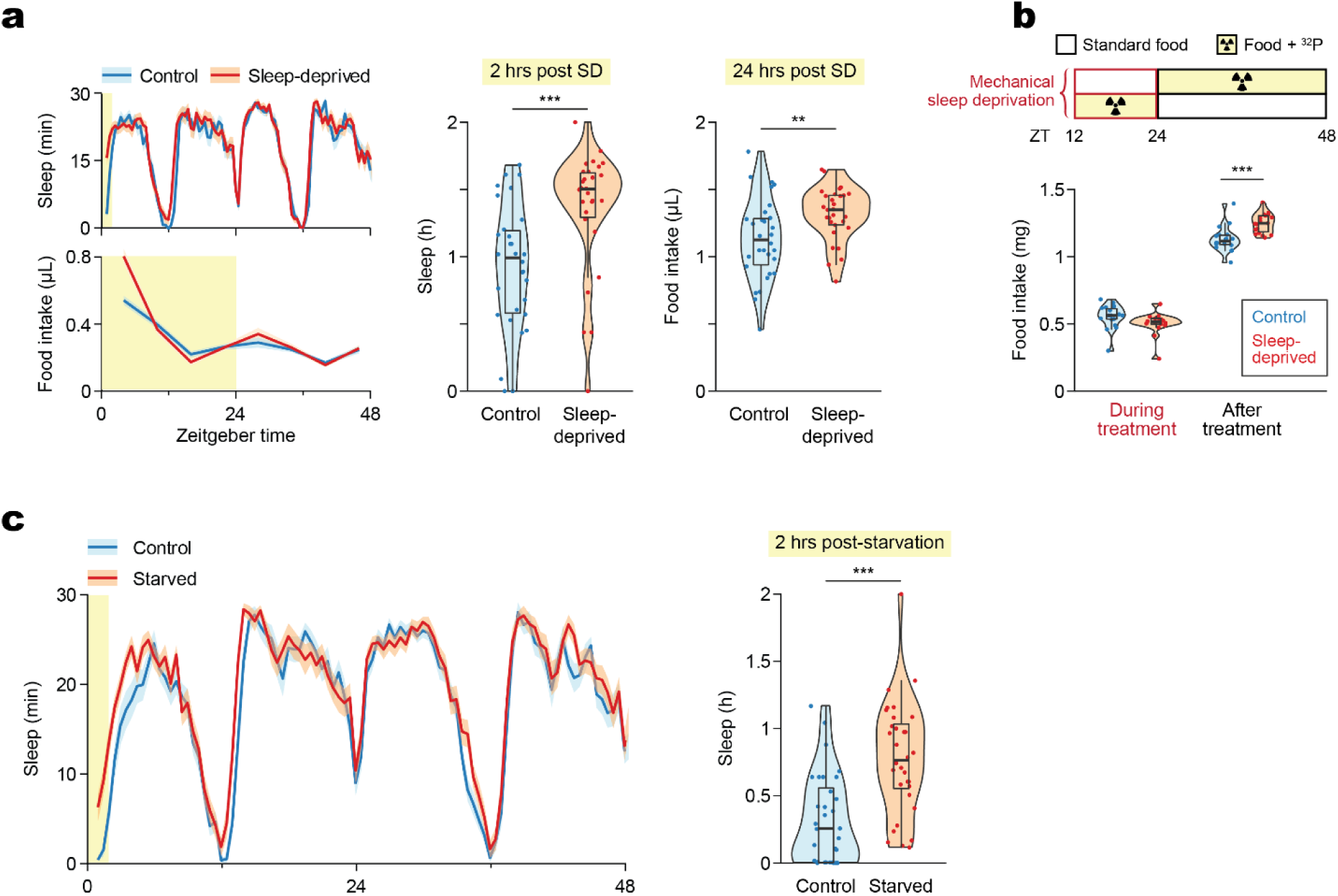
Mechanical disturbance and starvation suppress sleep and lead to post-deprivation sleep rebound and hyperphagia. (a) Mechanical sleep deprivation overnight leads to sleep rebound and hyperphagia the following day, measured in the Activity Recording CAFE (ARC). N = 30 Canton-S males per condition. (b) Food intake quantified by radiolabeling of the medium shows no difference in consumption during sleep deprivation, and confirms hyperphagia after SD. *N* = 18 Canton-S males per condition. (c) Sleep rebound following starvation, measured in the ARC. *N* = 30 Canton-S males per condition. Box plots represent median and interquartile range. Lines represent mean and the shaded regions SEM. Asterisks denote significant difference between the control and the sleep-deprived group by Welch’s *t* test (a,c) or Tukey multiple comparisons of means (b) (**, *p* < 0.01; ***, *p* < 0.001). SD, Sleep Deprivation.

Manipulating food intake reciprocally affected sleep. Depriving flies of food overnight increased not only food intake but also total sleep the following day (Fig. 1c), consistent with previous studies on the antihypnotic effects of starvation^3^. In summary, loss of either sleep or food intake led to an increase in both.

What is the relationship between sleep loss, rebound sleep, and food intake? Are homeostatic sleep rebound and hyperphagia controlled by the same mechanisms and, if not, can they be experimentally decoupled? Does sleep loss have direct orexigenic effects, or is the increased feeding a response to altered energy states following sleep deprivation? To explore these questions, we thermogenetically activated several broad groups of neurons that cause soft or hard sleep loss^20^, allowing us to dissect the components of sleep homeostasis and their potential effects on feeding.

TrpA1-mediated stimulation of cholinergic neurons suppressed sleep and led to sleep rebound (Fig. 2a-b), as previously reported^20^. Additionally, the compensatory sleep rebound was accompanied by post-SD hyperphagia (Fig. 2b). Sleep rebound and hyperphagia were observed in both sexes (Fig. S1a). In contrast, thermogenetic activation of octopaminergic (Fig. 2c) or dopaminergic (Fig. 2d) neurons induced sleep suppression without statistically significant changes in rebound sleep or feeding.

**Figure 2.**
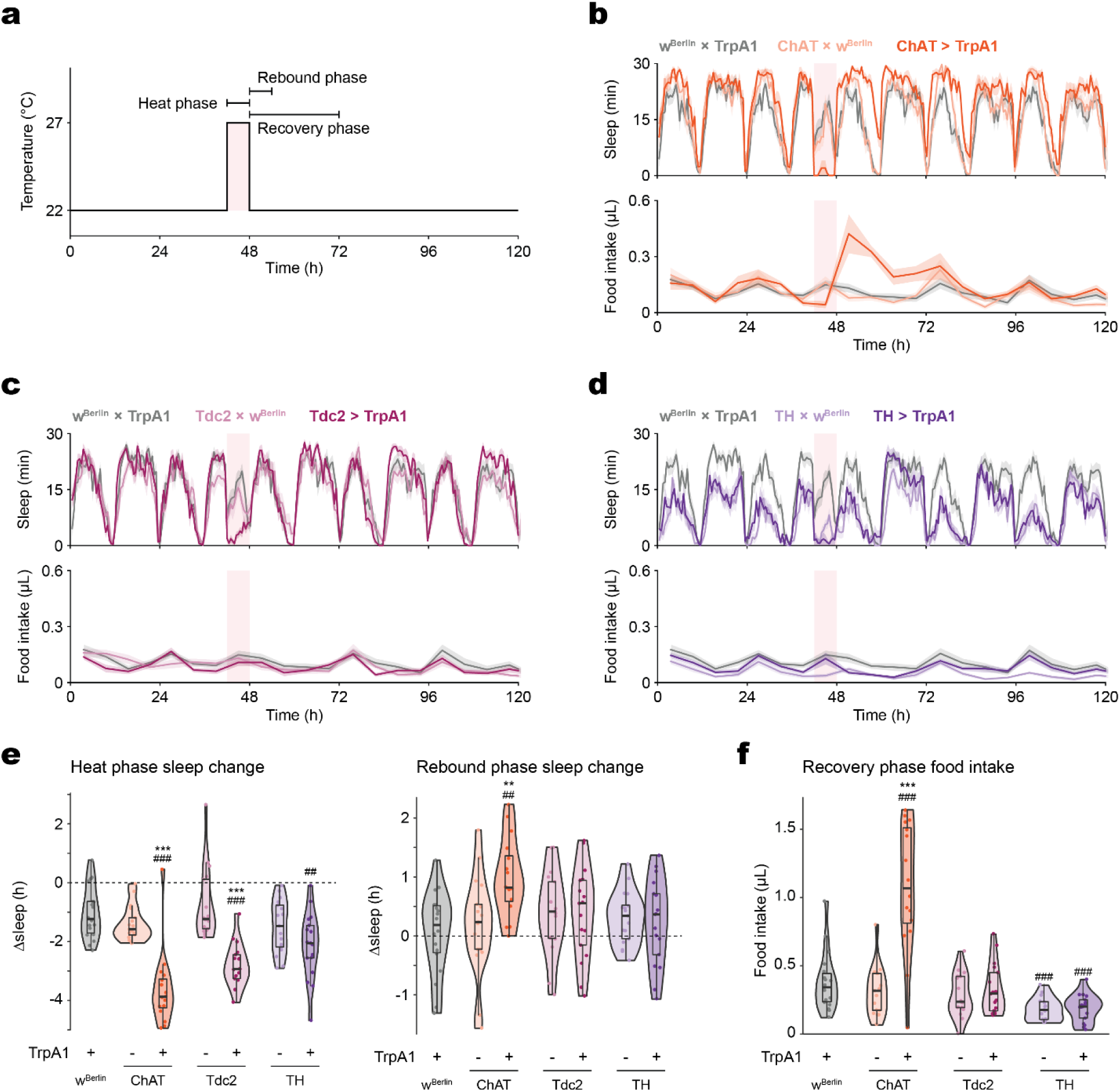
Food intake and sleep are altered during and following broad sleep-suppressing manipulations. (a) Schematic of the thermogenetic stimulation and references for the terms denoting sleep and feeding behaviors surrounding the stimulation. Baseline sleep is recorded at 22 °C prior to the stimulating heat phase, and thermogenetically stimulated (“heat phase”) behaviors are recorded at 27 °C unless specified otherwise. Heat and Rebound phase Δsleep are defined as the difference between total sleep during the heat (ZT 18-24) or rebound (ZT 0-6) phase and that of the same period during the baseline measurements from the previous day. Heat and Recovery phase feeding refer to total food intake during the 6 hours of the heat phase or the 24 hours following the heat stimulation, respectively. (b) Cholinergic stimulation elicits sleep loss and rebound, coupled with an increase in food consumption during the recovery phase. (c-d) Octopaminergic (c) and dopaminergic (d) stimulation elicit sleep loss but not sleep rebound or increased recovery feeding. (e) Stimulation of cholinergic, octopaminergic, or dopaminergic neurons suppress sleep, but only cholinergic stimulation is followed by sleep rebound. (f) Cholinergic stimulation results in increased recovery feeding. The light pink rectangular background indicates heat phase (b-d). Box plots represent median and interquartile range. All lines represent mean and the shaded regions SEM. *N* = 17-18 males per genotype. Following one-way ANOVA, significant differences were compared post-hoc using Tukey pairwise comparisons between the experimental and each of the control lines. Significant differences from the post-hoc comparisons are noted as: #, significantly different from TrpA1 control; *, significantly different from GAL4 control. The number of symbols denote the *p* value of the difference between the control and the sleep-deprived group (**/##, *p* < 0.01; ***/###, *p* < 0.001).

Cholinergic neuron stimulation, which induces rebound sleep, resulted in greater sleep loss than the octopaminergic and dopaminergic manipulations (Fig. 2e). To test whether the degree of sleep loss drives rebound sleep, a longer, 12-hour stimulation was tested. Extended activation of octopaminergic neurons further suppressed sleep but had no effect on sleep rebound or food consumption afterwards (Fig. S1b).

Dopaminergic neurons, when stimulated for longer, suppressed sleep and elicited sleep rebound, but had no effect on post-SD food intake (Fig. S1b). Instead, hyperphagia was observed during the treatment period itself. These results suggest that hard sleep loss is not only due to the degree of sleep loss, but that both a threshold level and ‘class’ of SD is necessary to trigger sleep rebound.

To better understand the relationship between sleep homeostasis and feeding behavior, we examined the effects of activating more restricted subsets of cholinergic neuronal populations implicated in various aspects of sleep regulation: sleep homeostasis; arousal (suppresses sleep but does not elicit sleep rebound); or sleep pressure (increases sleep following neuron activation with no sleep loss during treatment)^19,20^.

Consistent with previous findings^20^, activation of neurons labeled by 24C10-GAL4 or the intersection of 24C10-GAL4 and ppk-GAL4 (24C10∩ppk) suppressed sleep and elicited sleep rebound (Fig. 3a, Fig. S2a). There was no post-SD hyperphagia, but food intake was elevated during neuron stimulation, similar to the feeding phenotype that accompanied the longer activation of dopaminergic neurons (Fig. S1b). Activation of neurons labeled by 44F01- or 60D04-GAL4 suppressed sleep during the heat phase but did not evoke homeostatic sleep rebound (Fig. 3b), consistent with previous work suggesting that these neurons are directly wake-promoting and bypass the sleep homeostat^20^. The 44F01 and 60D04 manipulations did not affect food intake during or after activation (Fig. 3b).

**Figure 3.**
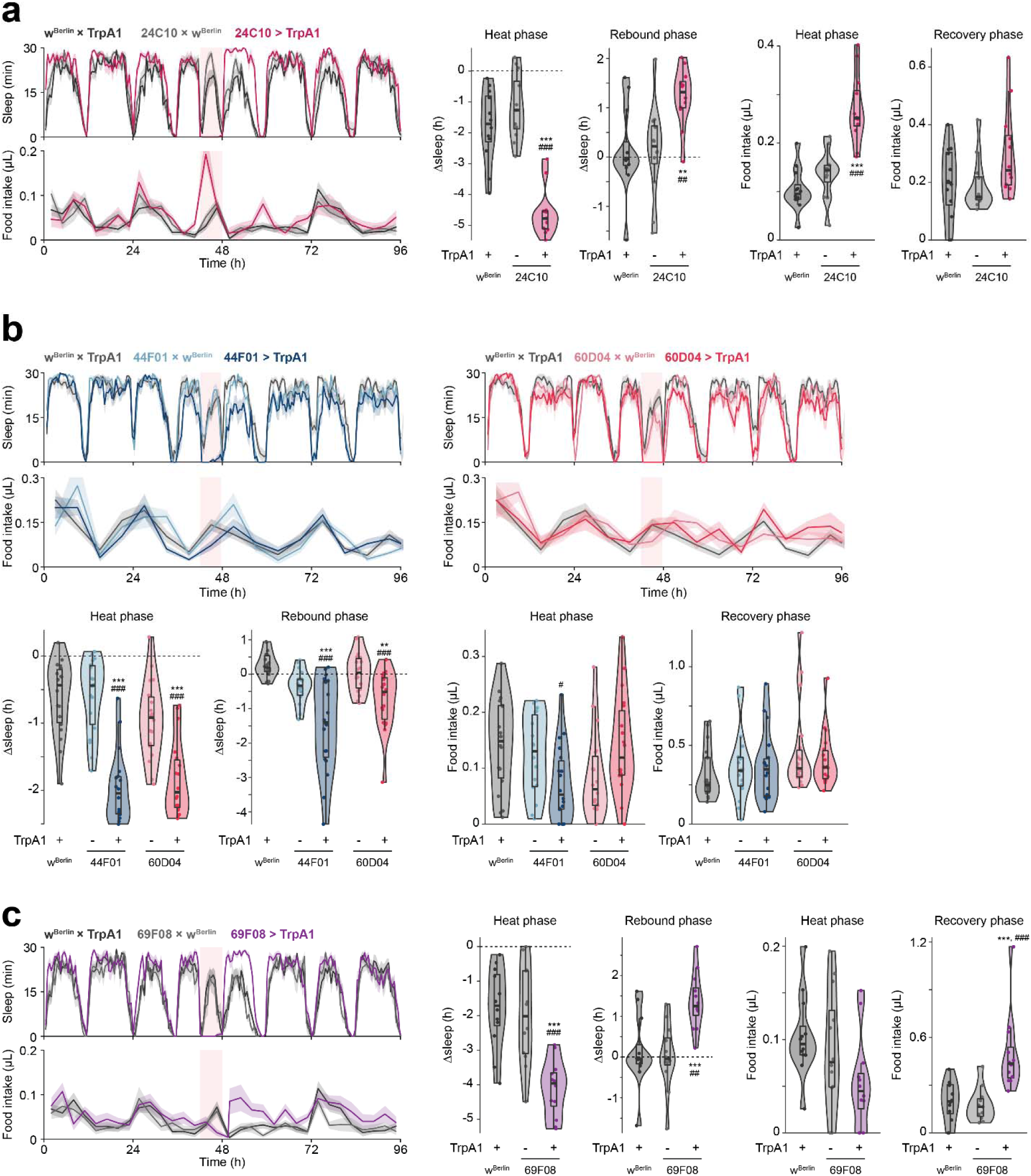
Thermogenetic stimulation of various subsets of cholinergic neurons causes soft or hard sleep loss. (a) 24C10-neuron stimulation elicits sleep loss and increased feeding during stimulation, as well as increased sleep during recovery phase. *N* = 12 males per genotype. (b) Stimulation of 44F01 or 60D04 neurons suppresses sleep without post-SD hypersomnia or hyperphagia. *N* = 6 males per genotype. (c) Stimulation of R2 neurons using 69F08-GAL4 driver suppresses sleep and food intake during the stimulation, with increased post-SD sleep rebound and food intake. *N* = 12 males per genotype. Box plots represent median and interquartile range. All lines represent mean and the shaded regions SEM. Following one-way ANOVA, significant differences were compared post-hoc using Games-Howell (if variance not homogeneous by Levene’s test) or Tukey pairwise comparisons (if variance homogeneous) between the experimental and each of the control lines. Significant differences from the post-hoc comparisons are noted as: #, significantly different from TrpA1 control; *, significantly different from GAL4 control. The number of symbols denote the *p* value of the difference between the control and the sleep-deprived group (*/#, *p* < 0.05; **/##, *p* < 0.01; ***/###, *p* < 0.001).

Previous studies have found that stimulation of 69F08 neurons in the ellipsoid body accrues sleep pressure either following^23^ or bypassing^19^ sleep deprivation. In our studies, activation of 69F08 neurons suppressed sleep, and induced both sleep rebound and post-SD hyperphagia while leaving food intake during the heat phase unaffected (Fig. 3c). We confirmed the sleep loss and subsequent rebound with 69F08 activation in females as well (Fig. S2b). Additionally, 69F08 activation decreased arousal threshold (Fig. S2c), suggesting that 69F08 neurons are wake-promoting. The sleep suppression was not due to the upright placement of the behavioral chambers, since 69F08 stimulation similarly suppressed sleep in ARC chambers oriented horizontally, mimicking the orientation in the widely used Drosophila Activity Monitor (DAM; TriKinetics) system (Fig. S2d).

The DAM system deduces activity levels and sleep based on the number of times an animal crosses an infrared beam that bisects the chamber^24^. This method assumes that any change in activity around the chamber midline is proportional to movement in unsampled areas—i.e., both ends of the chamber. Thus, single-beam DAM systems cannot accurately reflect the activity of manipulations that alter the spatial distributions of the movements^25^. In contrast, the ARC tracks the planar location of the animal, providing a more accurate measure of activity. We hypothesized that the particular assay used might have affected the detection of movement and sleep in 69F08 neuron-stimulated animals, consistent with other reports of artifactual hyperactivity using the DAM system^25,26^.

To reconcile the discrepancy between our 69F08 activation results and the previously reported sleep phenotype^19^, we ran post hoc analyses on the movement characteristics of the subjects in our original experiment. We found that 69F08 activation altered the spatial distribution of fly movements, where the experimental animals spent more time taking shorter strides in the bottom half of the chambers during stimulation (Fig. S2e).

We also reanalyzed the activity in a DAM-like manner—i.e., counting the number of times the flies crossed an imaginary line bisecting the chamber. Not surprisingly, sleep estimated using this method failed to capture the sleep loss during 69F08-neuron stimulation (Fig. S2e). Collectively, these results support the conclusion that increased sleep following 69F08 neuron activation is indeed sleep rebound and is not independent of homeostatic sleep debt^19,23^.

We hypothesized that the association of hard sleep loss with increased feeding might be due to changes in metabolic demands or energy balance during sleep loss. Thus, we next measured CO_2_ production as a proxy for metabolic rate during various neuronal manipulations. Increased respiration at 27 °C, compared to that at 22 °C, was observed in all manipulations associated with hard sleep loss and increased feeding, as well as with the octopaminergic manipulation (Fig. 4a-d). CO_2_ production was not affected by stimulation of 44F01 and 60D04—manipulations that also did not induce sleep rebound or affect feeding behavior (Fig. 4e).

**Figure 4.**
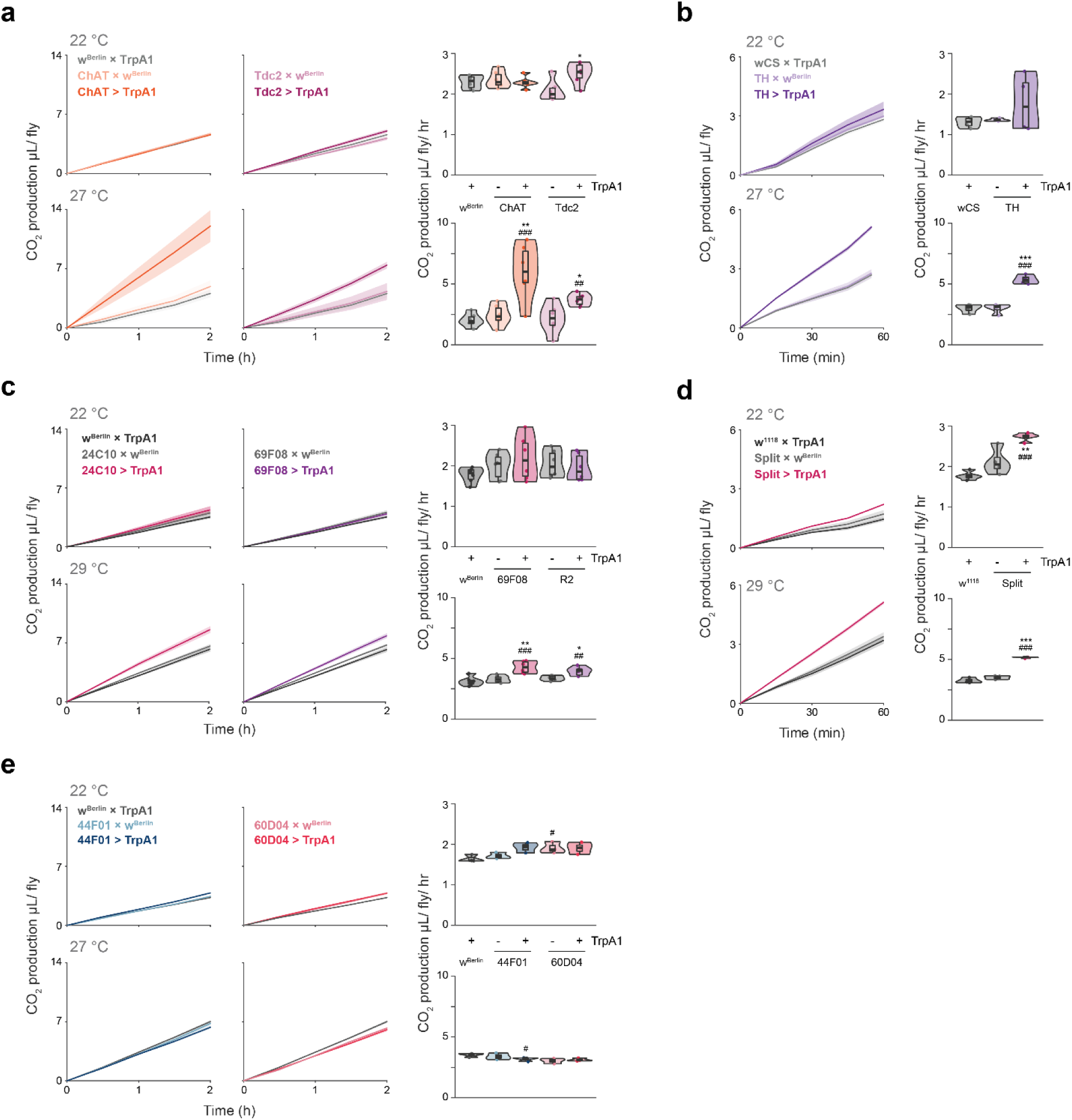
Increased respiratory metabolism is associated with manipulations that induce hard sleep loss. (a-e) 2-hour time-course (in 30-minute bins) or average CO_2_ production during stimulation of the noted neurons at the indicated temperatures. *N* = 6 cohorts per genotype, 4 males per cohort. Box plots represent median and interquartile range. Lines represent mean and the shaded regions SEM. Following one-way ANOVA, significant differences were checked for homogeneity of variance using Levene’s test and post-hoc comparisons between the experimental and each of the control lines were performed using Tukey’s multiple comparisons test. Significant differences from the post-hoc comparisons are noted as: #, significantly different from TrpA1 control; *, significantly different from GAL4 control. The number of symbols denote the *p* value of the difference between the control and the sleep-deprived group (*/#, *p* < 0.05; **/##, *p* < 0.01; ***/###, *p* < 0.001).

It is unclear whether hyperphagia, observed in all hard sleep loss manipulations, is merely a readout for energy deficit, or if it more directly influences sleep pressure. The former seems more likely, since manipulations that show increased food intake during stimulation, sufficient to eliminate post-treatment hunger, do not eliminate subsequent sleep rebound (Fig. 3a, Fig. S1b, Fig. S2a). Additionally, preventing food access during stimulation in these manipulations did not further increase sleep rebound compared to controls (Fig. 5). However, these studies are complicated by the possibility of a ceiling effect with the amount of sleep loss that can be induced. Interestingly, starvation during stimulation did not further increase recovery food intake compared to controls, suggesting that the manipulations tested (TH- and 24C10∩ppk-neuron stimulation) do not directly affect homeostatic feeding (Fig. 5). Taken together, our results suggest that hard sleep loss and energy deficit drive sleep and feeding pressures concomitantly.

**Figure 5.**
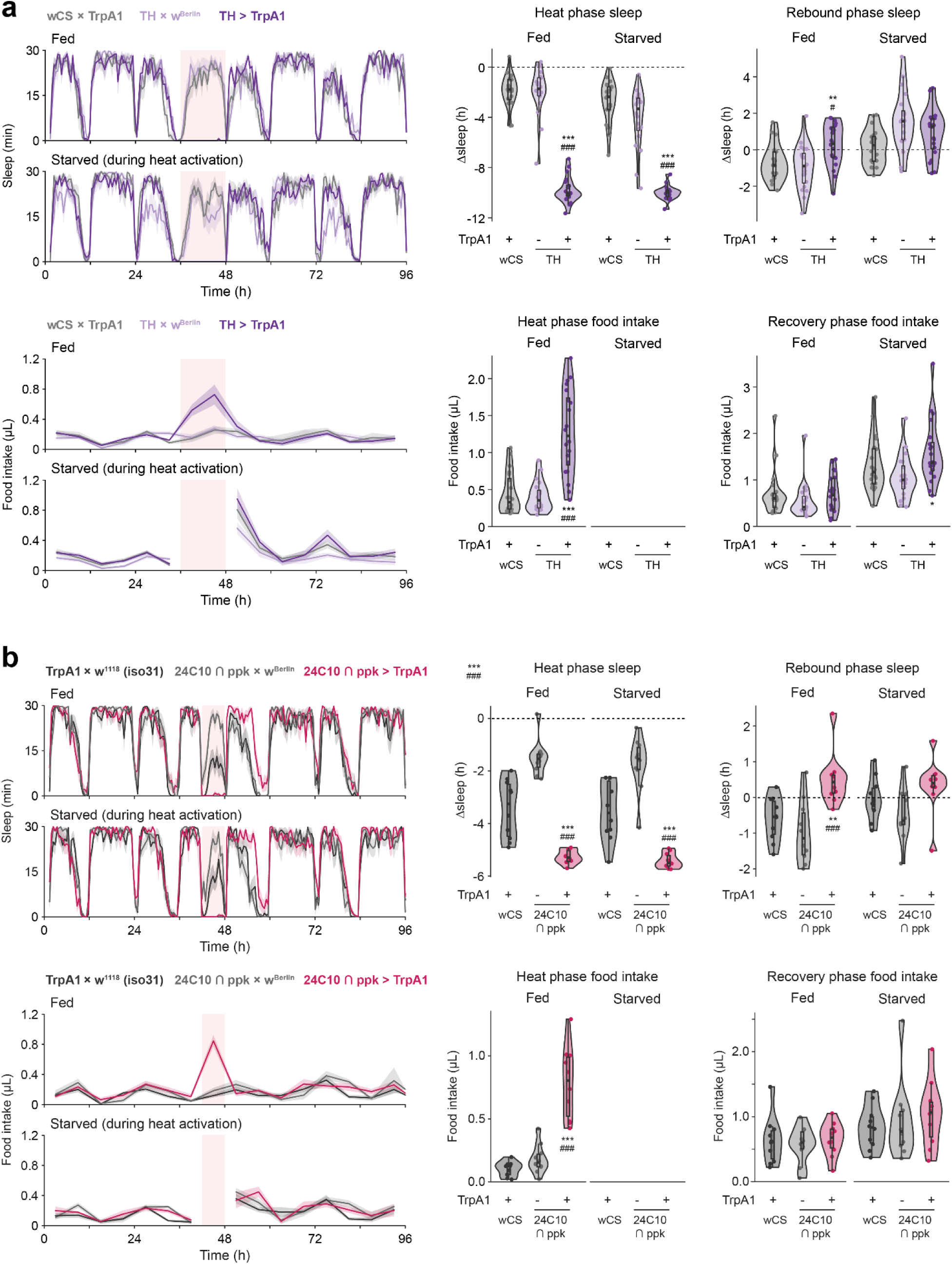
Starvation during neuronal stimulations that accompany increased food intake does not increase rebound sleep or recovery food intake. (a,b) Prohibiting access to food during the 12-hour TH-(a) or 6-hr 24C10∩ppk-(b) stimulation-induced SD is insufficient to increase sleep rebound or recovery food intake. Following one-way ANOVA, post-hoc Tukey pairwise comparisons were performed between the experimental (GAL4 > TrpA1) and each of the control lines. Significant differences from the post-hoc comparisons are noted as: #, significantly different from TrpA1 control (wCS × TrpA1); *, significantly different from GAL4 control (GAL4 × w^Berlin^). *N* = 10 males per genotype and treatment.

## Discussion

Our findings expand upon previous work demonstrating that sleep deprivation increases metabolism^21^. While increased metabolic rates have been observed in sleep-deprived animals^27,28^, our studies suggest negative energy balance is a critical factor in driving rebound sleep. However, we tested a limited set of sleep-suppressing manipulations. Assessing how other sleep-altering manipulations affect metabolism and feeding behavior could further support our interpretation.

Although hyperphagia is observed in all instances of hard sleep loss, its variable timing (during or after SD), and the lack of an additive effect on sleep rebound with starvation and wake-promoting neuron stimulation (Fig. 5), suggest that feeding pressure is not directly driving sleep pressure. Instead, it may be merely a readout of energy deficit.

This may also explain why Tdc2 stimulation increases respiration without inducing sleep rebound—no hyperphagia was observed, suggesting that these flies were not in energy deficit. Since CO_2_ production alone does not always accurately reflect energy expenditure^29^, it is possible that additional metabolic measurements may reveal a more consistent factor that ties energy balance to sleep rebound. It is also possible that Tdc2 stimulation suppresses the feeding response to metabolic expenditure. Octopaminergic stimulation has been shown to suppress sleep rebound following mechanical sleep deprivation, and octopaminergic MS1 neurons suppress sleep in favor of mating^20,30^.

Sleep is critical for the normal day-to-day function of most animals and, in flies, deficient or irregular sleep is associated with numerous adverse effects on physiology and metabolism^31–33^. Sleep homeostasis and rebound sleep serve restorative functions, perhaps only upon signaling from sleep debt. Our findings suggest that one such signal might be negative energy balance. This idea is consistent with the hypothesis that one of the functions of sleep is to conserve energy, especially since sleep typically decreases metabolic rate^21,34,35^, and supports previous studies implicating the role of energy-sensing genes in sleep homeostasis^11,36,37^.

While our study provides compelling evidence for the interplay of energy balance and sleep homeostasis, the use of *Drosophila melanogaster*, which enables access to powerful genetic manipulations, requires careful consideration when extrapolating findings to mammals with more complex metabolic systems. Additionally, it remains possible that the link between energy deficit and sleep rebound involves not only direct metabolic sensing, but also potential secondary stress responses or circadian misalignment induced by sleep deprivation. Further research using complementary model systems and investigating additional aspects of metabolism could provide a more complete understanding of sleep regulation.

In conclusion, our study reveals a link between energy balance and mechanisms of sleep homeostasis. This finding challenges the traditional view of sleep regulation as being primarily driven by the accumulation of sleep debt and suggests a more integrative model where sleep serves a critical role in metabolic restoration. Our work highlights the potential for therapeutic approaches to sleep disorders or metabolic conditions through the manipulation of energy states. Future studies exploring the specific metabolic signals communicating with sleep homeostatic centers will further advance our understanding of this fundamental relationship.

## Materials and methods

### Fly lines and husbandry

Flies were reared and maintained on a standard cornmeal-sucrose-yeast medium (3.1% active dry yeast, 0.7% agar, 5.8% cornmeal, 1.2% sucrose (all w/v), 1% propionic acid (v/v) and 0.22% Tegosept (w/v, pre-dissolved in ethanol)) in a light-, temperature-, and humidity-controlled incubator (12-/12-h light/dark cycle, 25°C, 60% relative humidity), unless otherwise specified. Young adults (2-4 days old) were separated into single-sex vials (10-15 flies/vial) under CO_2_ anesthesia and transferred to fresh food every other day until 7-10 days old. Female flies collected this way are considered once-mated. Transgenic flies were typically outcrossed for >10 generations to the indicated strain (a “Cantonized” white-eyed line, wCS; or white-eyed Berlin, w^Berlin^) that was confirmed to be free of *Wolbachia* by PCR analysis.

For thermogenetic experiments, flies were reared and maintained at 18 °C except during testing, during which they were transferred to control (22 °C) or experimental temperatures. 69F08- and ppk∩24C10-neuron stimulation was induced at 29 °C. All other thermogenetic stimulations were performed at 27 °C.

The following lines were acquired from Bloomington Drosophila Stock Center (BDSC): ChAT-GAL4 (#6793), TH-GAL4 (#8848), Tdc2-GAL4 (#9313), 24C10-GAL4 (#49075), 44F01-GAL4 (#45313), 60D04-GAL4 (#45356), and 69F08-GAL4 (#39499). The split-GAL4 constructs (ppk∩24C10) were acquired from Dr. William Joiner. UAS-TrpA1 was acquired from Dr. Ulrike Heberlein. All lines except the ppk∩24C10 split-GAL4 were outcrossed. Nonetheless, the appropriate hybrid backgrounds were used as controls for all comparisons.

### Diets

Liquid diets were filtered (0.22-μm cellulose acetate sterile syringe filter, VWR). Solid diets were prepared with Bacto™ Agar (BD Diagnostics) and a propionic/phosphoric acid mix was added to prevent bacterial growth^38^. ARC experiments were conducted with a standard liquid diet of 5% sucrose + 5% yeast extract (both w/v) except during starvation, during which the feeding capillaries were filled with deionized H_2_O.

### Mechanical sleep deprivation

To mechanically prevent flies from sleeping, we used a custom-built device that agitates fly vials in a 3-dimensional space. The device consisted of a 3D-printed vial carrier secured to two stepper motors (Zhengke Motor #ZGA28RP37.9i) controlled by an Arduino microcontroller. One motor rotated the carrier in a cartwheeling motion and the other delivered short, timed punches to further dislodge and disrupt the flies (Supplemental Video). This agitation cycle was repeated every 5 minutes to effectively prevent sleep. Sleep deprivation with this device was confirmed by observing subsequent sleep rebound compared to undisturbed controls.

### Sleep and feeding behavior measurement (ARC)

We used the Activity Recording CAFE (ARC) system^22^, a modified version of CApillary FEeder (CAFE)^39^, to simultaneously measure sleep and food intake. Flies were individually loaded by mouth pipette into chambers with access to liquid food (5% sucrose + 5% yeast extract, both w/v) in a 5-µL glass capillary. Prior to recording, flies were acclimated to the chambers overnight. Feeding events were detected as sudden, suprathreshold drops in the liquid meniscus level. Arousal threshold was measured as previously described^4,22^. The analyses excluded animals that died during the experiment to avoid artificially high arousal threshold values from non-responsiveness. General malaise or other artifacts were monitored with real-time camera output as well as through post hoc analysis of the animal movements.

For the horizontally placed ARC experiment, the ARC chambers were loaded with an aliquot of agar on the bottom as in the typical setup. Afterwards, flies were anesthetized by CO_2_ and loaded into the chambers. Before the flies recovered from anesthesia, the top of the chamber was fitted with 200-μL unfiltered pipette tips filled with solid food (5% sucrose, 5% yeast extract, and 1% agar, all w/v), instead of capillary tubes holding liquid food. The narrow ends of the pipette tips were trimmed to allow access by flies, and the opposite ends were wrapped with Parafilm® to reduce evaporation. The chambers were laid flat or horizontally on the incubator shelf and cameras were mounted above. Animal movements were tracked and recorded as in the standard ARC setup, except that food intake was not monitored.

### Radiolabeled food intake

To quantify food intake during and following mechanical sleep deprivation, we measured consumption of radiolabeled solid food in standard vials^40,41^. Flies were divided into four experimental groups: sleep-deprived with radiolabeled (“hot”) food, sleep-deprived with non-radiolabeled (“cold”) food, undisturbed with hot food, and undisturbed with cold food. Flies in the hot food group had access to food mixed with 1–2 μCi/mL [α-^32^P] during the sleep deprivation period. Following the 12 hours of sleep deprivation, flies in the hot food groups were collected in empty vials and frozen at -80 °C. Flies from the cold food groups were subsequently allowed to feed *ad libitum* on hot food for 24 hours and then frozen in empty vials.

Accumulated radiotracer was quantified in whole flies by liquid scintillation^40,41^. Pre-weighed aliquots of non-solidified radiolabeled media were used as a conversion factor to determine food mass consumed. We have previously confirmed that the low concentration of radiolabel used (<1 nM α·^32^P-dCTP) does not affect feeding behavior^40^.

### Respiratory metabolic expenditure

We used CO_2_ generation as a proxy for overall energy expenditure^42^, using the previously described method with slight modifications^43^. A metabolic chamber was fabricated using a non-barrier P1000 micropipette tip (Fisher Scientific cat no. 02-681-172) containing soda lime (Ward’s Science #470302-416), separated from the residential area by a piece of low-density packing foam. The narrow end of the tip was fitted with a 50-µL glass capillary (Pyrex Disposable Micro-Sampling Pipets, Corning cat no. 7099S-50) and sealed with hot glue. A 100-µL droplet of 5% sucrose, 5% yeast extract, and 1% agar (all w/v) was applied to the wall of each metabolic chamber, including the blank control. Each metabolic chamber was loaded with 4-5 flies using a mouth aspirator and sealed with non-hardening modeling clay (Plastilina, Sargent Art #22-7688). The blank chambers, used as controls for atmospheric pressure, were constructed identically except no flies were loaded. A thin layer chromatography (TLC) box was filled with red-dyed water (Eosin Y, Fisher Chemical cat no. E511-100) and fitted with a 3D-printed rack that suspended the metabolic chambers at an equal height with all glass capillary tips below the surface of the dyed water. The prepared TLC box was sealed with high vacuum grease (Dow Corning) and placed in front of a camera in a temperature-controlled incubator.

PhenoCapture (www.phenocapture.com) software was used to capture a series of pictures of the TLC box at 5-minute intervals, and the timelapse images were analyzed using FiJi^44^ to automatically measure the meniscus level of the dyed water inside the glass capillaries. The changes in the meniscus levels were assumed to be directly proportional to respiratory metabolic expenditure.

### Data analysis

All analyses were performed using the R statistical package^45^. For one-factor experiments, normally distributed data were first checked for homogeneity of variances with Levene’s Test followed by standard analysis of variance (ANOVA) or, in case of unequal variance, Welch’s variant of ANOVA. Tukey’s multiple comparisons of means test was used for post hoc comparisons between groups and data that were not normally distributed were analyzed using the Kruskal-Wallis test. When comparing between phases of the same experiment, we used repeated measures ANOVA or, if the data were of unequal variances or not normally distributed, the Friedman test.

Significant interaction terms were followed up with post hoc within-genotype comparisons between phases with Bonferroni adjustments for multiple comparisons. An α level of .05 and power (1-β) of .80 were used for all statistical tests.

## Acknowledgments

We thank Dr. Binbin Wu for helpful comments on this manuscript. This work was funded by the NIH (R01DC020031, W.W.J.).

## Author Contributions

Conceptualization, Methodology, Data collection, and Data Analysis: S.J.P., K.R.M., and W.W.J.; Supervision: W.W.J.; Writing—original draft: S.J.P. and K.R.M.; Writing—review and editing: S.J.P., K.R.M., and W.W.J.

## Declaration of Interests

The authors declare no competing interests.

---

**Supplemental Figure S1.**
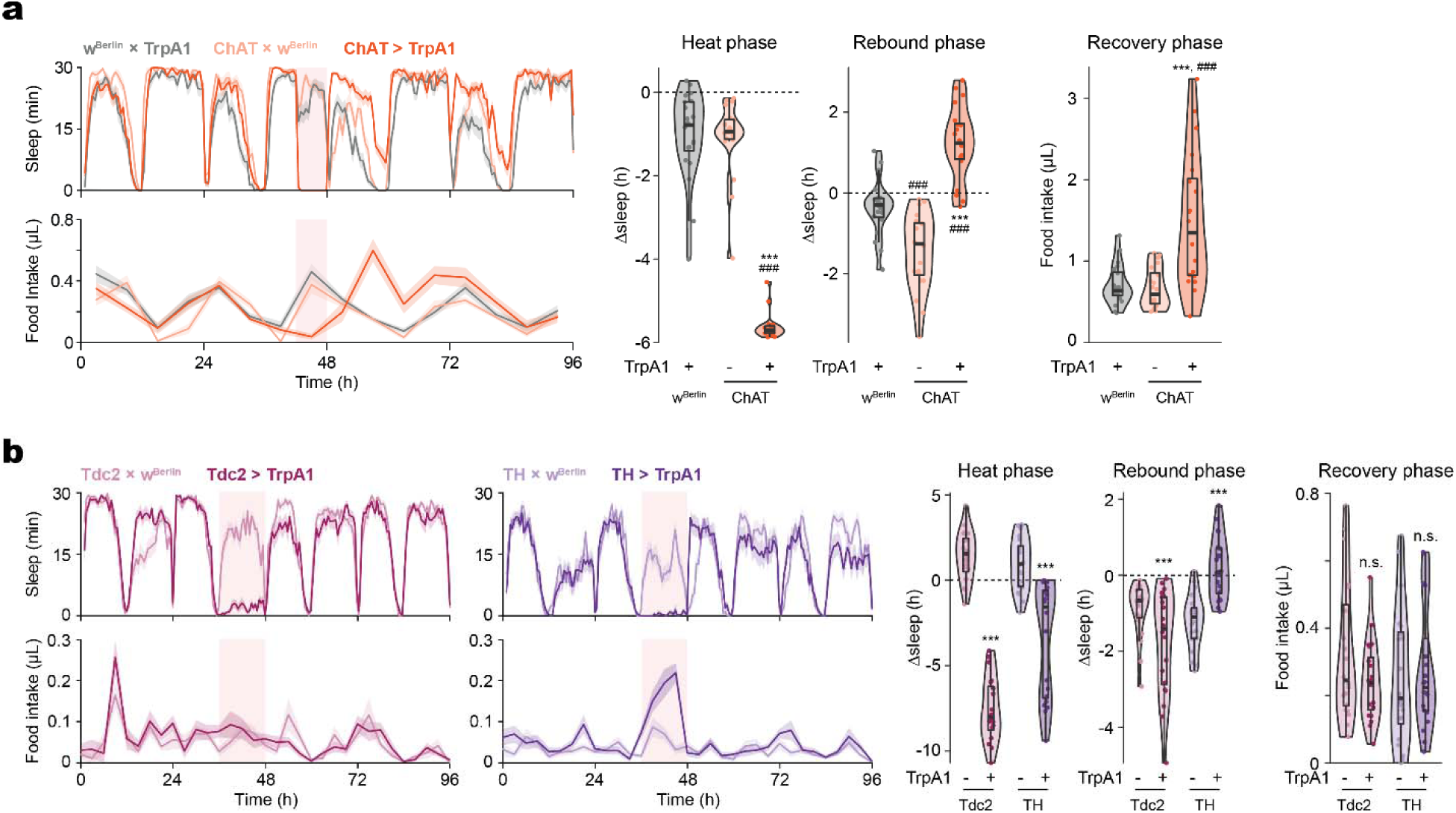
Food intake and sleep are altered during and following broad sleep-suppressing manipulations. (a) Cholinergic stimulation elicits sleep loss, sleep rebound, and post-SD overfeeding in females (*N* = 20 per genotype). (b) Prolonged (12 hr) stimulation of octopaminergic neurons suppresses sleep but does not affect post-SD sleep or food intake, whereas prolonged stimulation of dopaminergic neurons suppresses sleep, increases food intake during the heat phase, and induces rebound sleep. The light pink rectangular background indicates heat phase. Box plots represent median and interquartile range. All lines represent mean and the shaded regions SEM. *N* = 15 males per genotype. Following one-way ANOVA, significant differences were compared post-hoc using Games-Howell (if variance not homogeneous by Levene’s test) or Tukey pairwise comparisons (if variance homogeneous) between the experimental and each of the control lines. Significant differences from the post-hoc comparisons are noted as: #, significantly different from TrpA1 control; *, significantly different from GAL4 control. The number of symbols denote the *p* value of the difference between the control and the sleep-deprived group (***/###, *p* < 0.001).

**Supplemental Figure S2.**
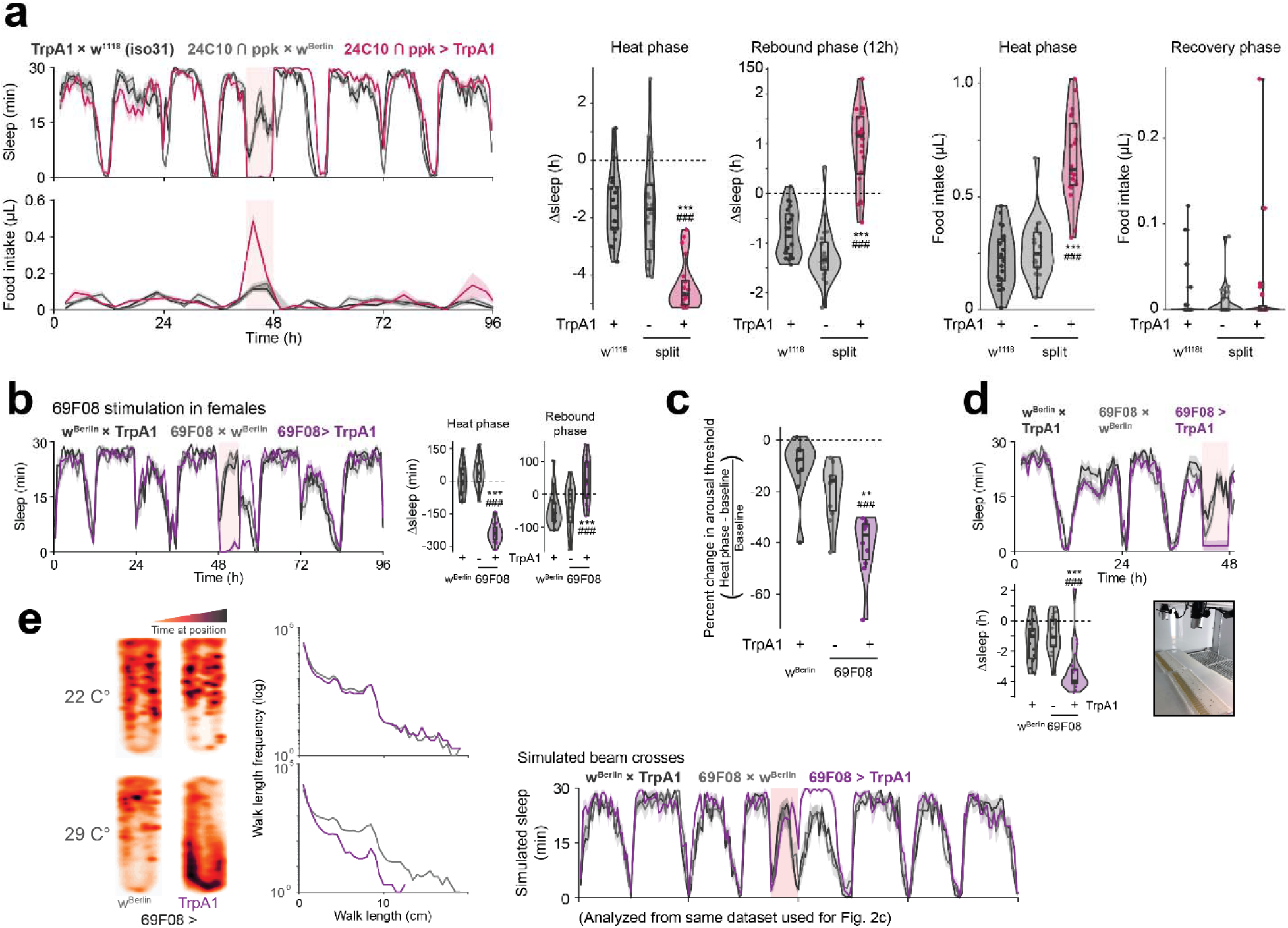
Thermogenetic stimulation of restricted subsets of cholinergic neurons causes hard sleep loss. (a) Stimulation of a more restricted subset of R2 neurons suppresses sleep and increases feeding during the stimulation, followed by sleep rebound. *N* = 20 males per genotype. (b) 69F08 stimulation also elicits sleep loss and rebound sleep in females. *N* = 20 females per genotype. (c) 69F08 stimulation decreases arousal threshold, consistent with sleep loss. *N* = 10 males per genotype. (d) Orienting the ARC chamber horizontally has no effect on the sleep loss observed during 69F08 stimulation, suggesting that sleep loss is not due to restrictions from the vertical orientation of the chamber. *N* = 20 males per genotype. (e) Spatial map (left) and walk length frequency histogram (middle) of animals within ARC chamber during 69F08 stimulation from Fig. 3c. Simulated bisecting beam crosses from the animal movements show unchanged sleep, contrary to the observation from animal tracking. Box plots represent median and interquartile range. Lines represent mean and the shaded regions SEM. Following one-way ANOVA, significant differences were checked for homogeneity of variance using Levene’s test and post-hoc comparisons between the experimental and each of the control lines were performed using Tukey’s multiple comparisons test. Significant differences from the post-hoc comparisons are noted as: #, significantly different from TrpA1 control; *, significantly different from GAL4 control. The number of symbols denote the *p* value of the difference between the control and the sleep-deprived group (*/#, *p* < 0.05; **/##, *p* < 0.01; ***/###, *p* < 0.001).

## Notes

### Competing Interest Statement

The authors have declared no competing interest.

### Summary of Updates

This version of the manuscript contains revised text and figures, updated references, and newly added acknowledgments

